# Carbon metabolism, transcriptome and RNA editome in developmental paths differentiation of *Coprinopsis cinerea*

**DOI:** 10.1101/819201

**Authors:** Yichun Xie, Jinhui Chang, Hoi Shan Kwan

**Author notes:** Address correspondence to Hoi Shan Kwan,.

## Abstract

The balance and interplay between sexual and asexual reproduction is one of the most attractive mysteries in fungi. The choice of developmental strategy reflects the ability of fungi to adapt to the changing environment. However, the evolution of developmental paths and the metabolic regulation during differentiation and morphogenesis are poorly understood. Here, we monitor the carbohydrate metabolism and gene expression regulation during the early differentiation process from the “fungal stem cell”, vegetative mycelium, to the highly differentiated tissue/cells, fruiting body, oidia or sclerotia, of a homokaryotic fruiting *Coprinopsis cinerea* strain A43mut B43mut pab1-1 #326, uncovering the systematic changes during morphogenesis and the evolutionary process of developmental strategies. Conversion between glucose and glycogen and conversion between glucose and beta-glucan are the main carbon flows in the differentiation processes. Genes related to carbohydrate transport and metabolism are significantly differentially expressed among paths. RNA editing, a novel layer of gene expression regulation, occurs in all four developmental paths and enriched in cytoskeleton and carbohydrate metabolic genes. It is developmentally regulated and evolutionarily conserved in basidiomycetes. Evolutionary transcriptomic analysis on four developmental paths showed that all transcriptomes are under purifying selection, and the more stressful the environment, the younger the transcriptome age. Oidiation has the lowest value of transcriptome age index (TAI) and transcriptome divergence index (TDI), while fruiting process has the highest of both indexes. These findings provide new insight to the regulations of carbon metabolism and gene expressions during fungal developmental paths differentiation.

**Importance:** Fungi is a group of species with high diversity and plays essential roles to the ecosystem. The life cycle of fungi is complex in structure and delicate in function. Choice of developmental strategies and internal changes within the organism are both important for the fungus to fulfill their ecological functions, reflecting the relationship between environment and the population. This study put the developmental process of vegetative growth, sexual and asexual reproduction, resistant structure formation of a classical model basidiomycetes fungus, *C. cinerea*, together for the first time to view the developmental paths differentiation process with physiology, transcriptomics and evolutionary prospects. Carbohydrate assays and RNA-seq showed the changes of the fungus. Our results fill the gaps on gene expression regulation during the early stage of developmental paths differentiation, and expand our understanding of the evolutionary process of life history and reproductive strategy in fungi.

## Introduction

The life cycle of fungi is one of the most delicate and mysterious development blueprints in the living world. Basidiomycetes, a group of advanced fungi, could diverge themselves into different developmental paths under specific environment (1–3). Triggering by the changes of environmental and internal physiological conditions, organism undergoes particular path and reaches the developmental destiny of forming asexual spores (oidia), sexual spores (basidia), monocellular or multicellular resting structures (chlamydospores and sclerotia) (4–7).

*Coprinopsis cinerea* is a model basidiomycete fungus which has multiple developmental paths and has well annotated genome of strain Okayama-7 #130 and strain A43mut B43mut pab1-1 #326 released (8, 9). *C. cinerea* can form several types of specialized reproductive structures to disperse and survive, according to the environmental conditions. Beginning from vegetative mycelium, the fungus goes along the developmental path of fruiting, oidiation or sclerotia formation (Fig. 1). During the differentiation process, several developmental changes occur in the organism, including regulations on gene expression, redistribution of chemicals, and specification on morphology (10–12). Fruiting body is a highly differentiated multicellular structure which forms during the sexual reproductive process (9). When the fungus is under nutrient depletion and exposing to low temperature and light-dark cycle, vegetative mycelium would undergo sexual reproduction. Hyphal knots develop from the aggregation of hyphae, they differentiate into fruiting body primordium, and finally become the mature fruiting body. During the maturation, basidiospores are produced by basidia in the cap and released as the cap autolyze. Oidia are asexual spores that form in favorable environment with light. *C. cinerea* can produce tremendous amount of oidia in a day. However, oidia are short-lived and fragile in stressful environment (13). Sclerotia are persistent resting structures that developed under continuous dark (14). They are multicellular structures in round shape, with internal medulla tissue and external rind tissue (15). The whole sclerotia are pigmented and in brown color (16).

**Fig 1.**
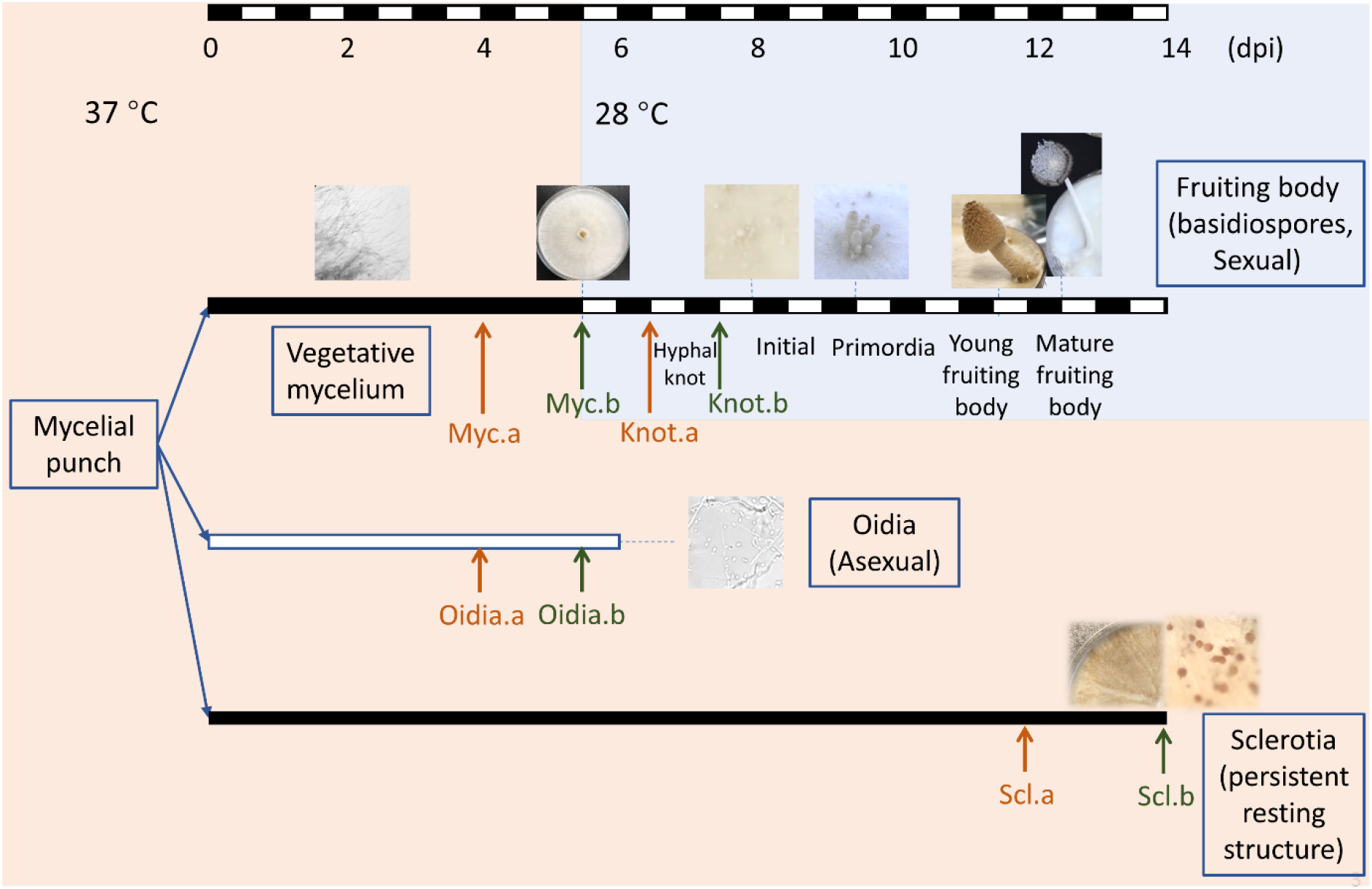
Developmental paths of *Coprinopsis cinerea* and experimental design. Mycelial punch was inoculated to each cellophane covered YMG plate and then incubated under different environment. Time scale bars indicate days post inoculation (dpi) and light exposure (white) or dark incubation (black) of each developmental process. Arrows indicate sample collection of time point a (before structure formation, developing colony, orange) and b (structure formed, developed colony, green).

In fungi, carbon metabolism is important to the differentiation process and complex morphogenesis. The transition from vegetative mycelia to each developmental destiny requires the core role of carbohydrate metabolism (17). Carbon metabolic flux describes the turn-over rate of molecules in carbon metabolic pathways. Such flux is regulated by enzymes and metabolic flux analysis (MFA) is important for metabolic adaptation studies (18–20). Through the carbon metabolic flux, materials and energy exchange between the organism and the environment, and they transfer within the organism and converse into different forms. Moreover, the construction of complex multicellular structure requires the conversion among different forms of carbohydrates (17, 21, 22). In the vegetative growth stage, mycelia intake simple sugar and synthesize glycogen (23–25). During the fruiting bodies or sclerotia formation, glycogen is transported from mycelium to newly-formed structures. It is broken down into glucose, and depleted along the maturation (25, 26). Beta-glucan is reserved in aging multicellular structures, such as fruiting bodies and sclerotia (5, 27, 28). Enzymes involved in fungal cell wall remodeling are up-regulated during the fruiting process (29–31). Despite carbohydrate metabolism has essential roles on differentiation and morphogenesis, how the transition is induced remains unclear.

RNA editing is regarded as a novel layer in gene expression regulation of fruiting body development (32). It is a kind of co-/post-transcriptional modification on RNA sequence, which can rewrite the genetic information in DNA at RNA level (33). By recoding the RNA sequence, RNA editing gives higher flexibility and more probability to transcriptome and proteome (34). Recently, several studies have been done in fungi, focusing on stage-specific and substrate-specific RNA editing events. Three ascomycetes, *Fusarium graminearum*, *Neurospora crassa* and *Neurospora tetrasperma*, have conserved stage-specific A-to-I RNA editing events during sexual reproduction (35). Developers of FairBase summarized the A-to-I RNA editing events in fungi, and emphasized that all these events are stage-specific and related to sexual reproduction (36). However, all 6 species included in FairBase are ascomycetes, and the A-to-I RNA editing preference in reported basidiomycetes are weak. Among the basidiomycetes, *Ganoderma lucidum* has stage-specific RNA editing events during fruiting body formation and 5 brown rot wood-decay fungi have substrate-specific RNA editing events (37, 38). These 6 species possess all types of RNA editing and with site densities much lower than those ascomycetes case studies.

The four developmental paths, namely vegetative growth, fruiting, oidiation and sclerotia formation, make *C. cinerea* able to adapt to divergent environment. Although the fruiting process have been studied and well described with transcriptomic and proteomic methods, our knowledge on oidia and sclerotia formation remains on single gene level (9, 39–42). The strain #326 used in this study is a homokaryotic strain with mutations in both mating type factor A and B (43). It shows the special feature of clamp formation and fruiting without mating, and possesses the ability of oidia and sclerotia formation, which gives us a chance to investigate developmental paths differentiation in *C. cinerea* with a clear genetic background. For the first time, we i) described carbon metabolic flux of four developmental paths; ii) figured out carbohydrate metabolism and energy production and conversion genes that are intensively regulated during morphogenesis; iii) clarified the evolutionary features of transcriptome profiles and explained the origin of developmental paths and adaptation to the environment; iv) uncovered the RNA editome in developmental paths differentiation. Our results enable us to have better understanding on the origin and regulation of developmental paths differentiation in advanced fungi.

## Results

### Temperature and light affect the divergence of developmental paths

To investigate the effect of environmental conditions on diverging the developmental paths, *C. cinerea* A43mut B43mut pab1-1 #326 cultures were treated with different combinations of temperature and light (Fig. 1). A 5 mm diameter mycelial punch was inoculated to each cellophane covered YMG agar plate (0.4% yeast extract, 1% malt extract, 0.4% glucose and 1.5% agar, 36 g medium per 90 mm diameter petri dish). The 5.5 days incubation in 37 °C with continuous dark allowed fully growth of vegetative mycelium. Fruiting body initials and mature fruiting bodies were seen on cultures after exposing to three and eight 12 h:12 h light-dark cycles in 28 °C, respectively. Oidia formation was highly induced under continuous light compared to continuous dark in 37 °C (P < 0.001, *N* = 6, Fig. S1a), with approx. 10^9^ oidia per plate on day 5.5. Mycelial growth rate had no significant difference between two conditions (P > 0.05, Fig. S1b). Sclerotia were firstly observed on day 14, after the undisturbed incubation in 37 °C with continuous dark. On day 21, 4.37 ± 0.69 × 10^4^ sclerotia were found per plate.

### Carbon metabolic flux differs among developmental paths

To evaluate the carbon metabolic flux in *C. cinerea*, cultures of four developmental paths were sampled on two time points, developing colonies (marked as a) or developed colonies (marked as b), with 5 biological replicates (Fig. 1). Neither hyphal knot nor sclerotia can be found on the developing colonies (time point a). Each sugar content was determined using chemical assays and measured by dry weight. Different types of carbohydrate were accumulated along the developmental paths (Fig. 2 and Table S1). During the vegetative growth process, glycogen was strongly accumulated to approx. 400 mg/g (dry weight, same below) in the hyphae and broken down when the colony was matured and shifted to the reproductive growth, reflecting the storage function of glycogen in fungal development. For vegetative mycelium, glucose content increased as colony grown and reached the peak of 266.16 ± 33.34 mg/g in fully grown mycelium. Mature colonies contain 1.5 times more beta-glucan than the growing ones. On the contrary, during oidiation, glucose content remains relatively constant at below 200 mg/g. Beta-glucan content was 100 ∼ 150 mg/g less in oidia than vegetative mycelium. Moreover, the amount of total sugar was also less in oidia-forming colonies (60-70 % of dry weight) than vegetative growth colonies (75-85 % of dry weight), suggesting that more non-sugar compounds, such as proteins and lipids, are produced during oidiation. Sugars, proteins and lipids that are synthesized in hyphae could be transferred and stored in oidia, preparing for the rapid germination in the surrounding favorable sediment. As nutrients in the sediment deplete, the colony turns to develop sexual reproductive structures or persistent resting structures according to the temperature and illumination. Being induced by low temperature and 12 h:12 h light-dark cycle, colonies with fully grown mycelia entered the sexual reproductive path and formed hyphal knots. During the transition, glucose content dropped down while glycogen content slightly increased. Beta-glucan took up more than one third of dry weight in the fruiting colony. When the colony is trapped in high temperature and dark environment, sclerotia are developed as persistent resting structure. Compare to vegetative mycelium, contents of monosaccharide, disaccharide and glycogen continuously dropped down in sclerotia, beta-glucan accumulated and took up over 50 % of the dry weight. In sclerotia, beta-glucan not only function as structuring constituent, but also the main type for carbon storage.

**Fig 2.**
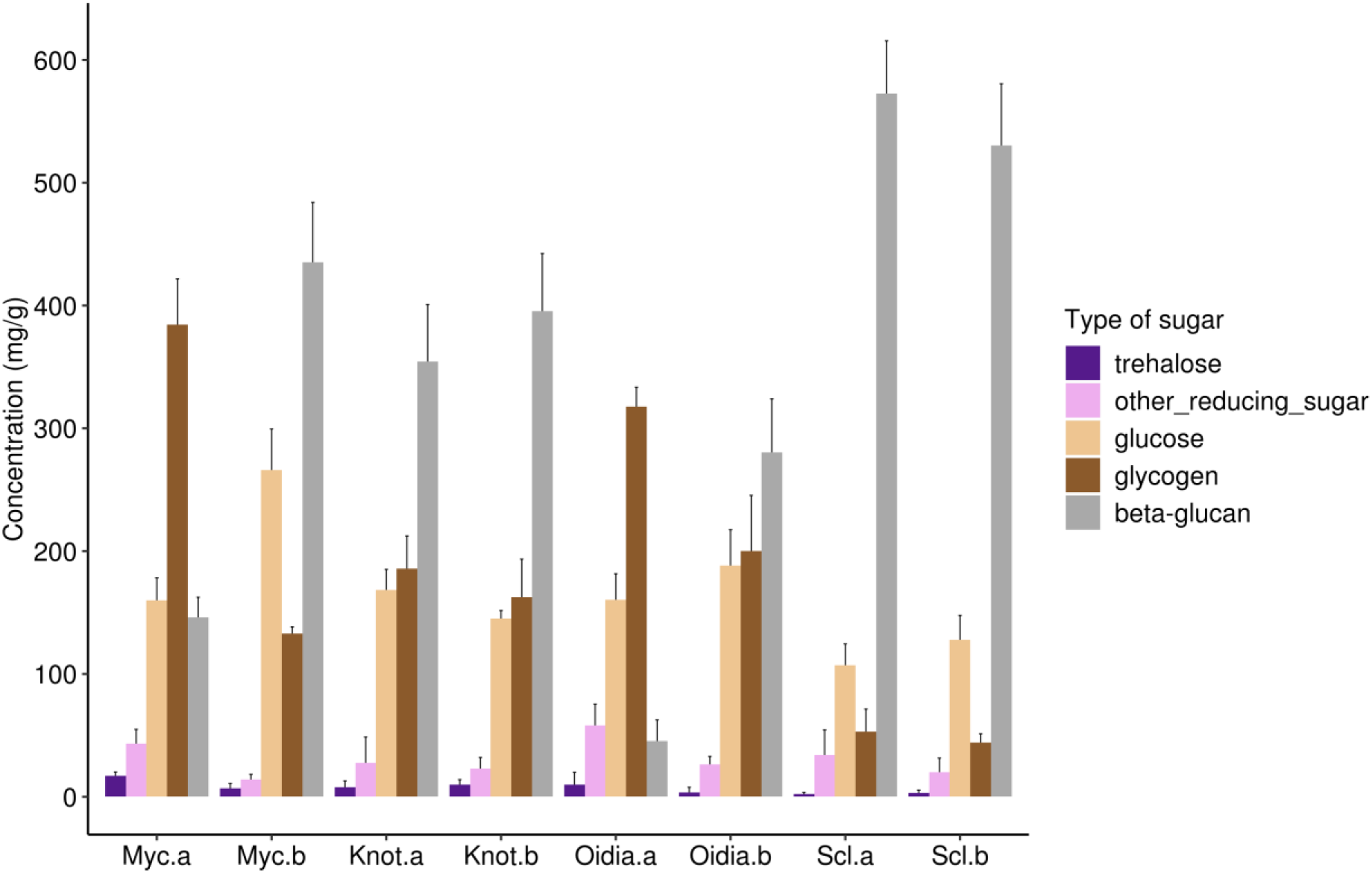
Carbohydrate content in different developmental paths. Error bar showing the standard deviation of 5 biological replicates. Statistic results present in Table S1.

### Transcriptome analysis on developmental paths differentiation

To figure out the black box behind the carbon metabolic flux in developmental paths differentiation, we studied the transcriptome of oidia and sclerotia formation process in *C. cinerea* for the first time, together with the fruiting process. Samples of developing colonies (time point a) were selected for RNA extraction, and RNA-seq was performed in biological triplicates. Sample and data quality were listed in supplementary method (text S1). Pearson’s correlation r^2^ value for replicates were between 0.9615 to 0.9883 (Fig. 3a), and Spearman’s correlation ρ^2^ value were range from 0.9661 to 0.9841 (Fig. 3b). Among these developmental paths, oidia showed the highest similarity to mycelium, and other paths were strongly different from each other.

**Fig 3.**
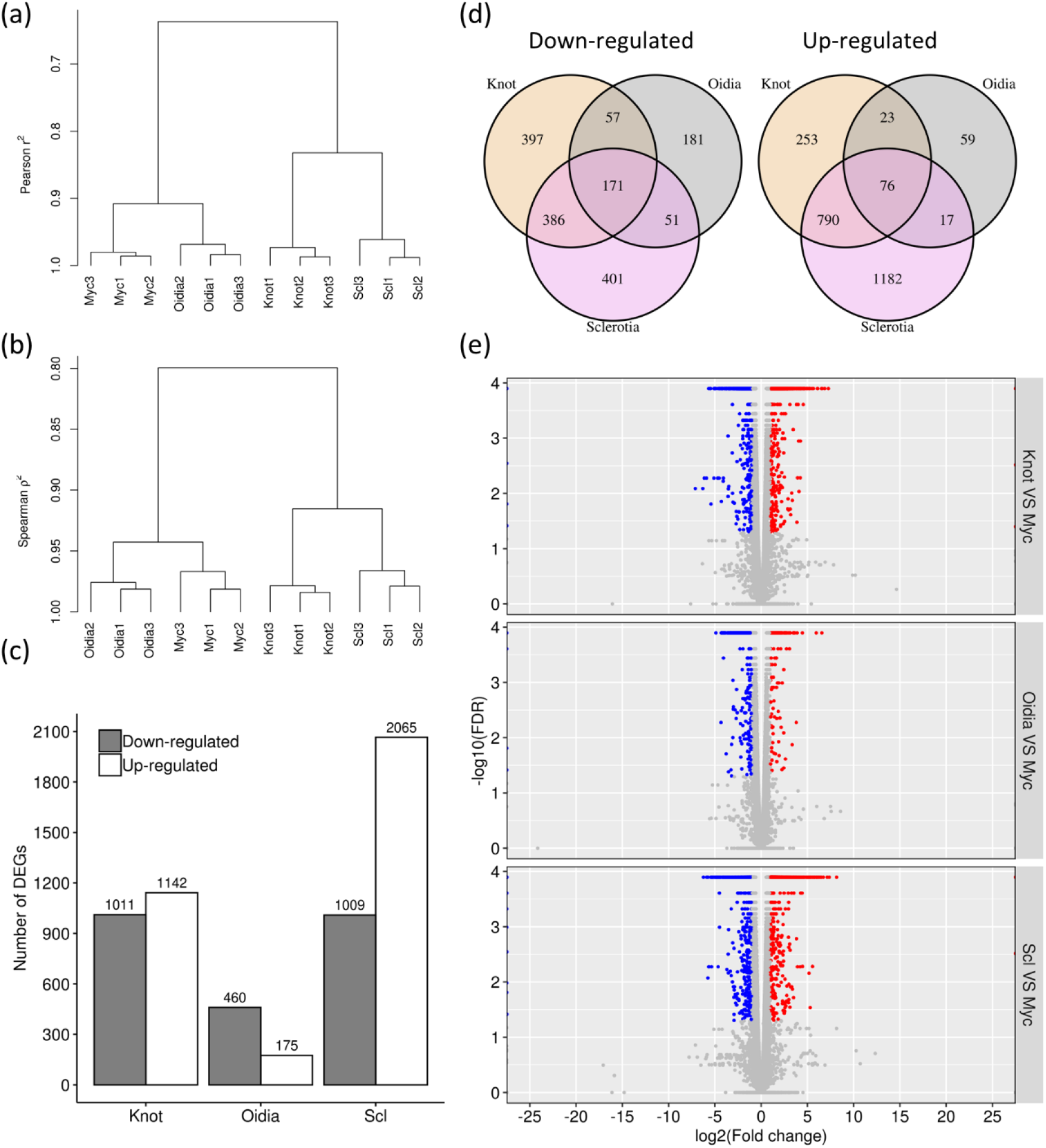
Transcriptome profiles of four developmental paths. (a-b) Hierarchical clustering of RNA samples using FPKM values with Pearson’s correlation (a) and Spearman’s correlation (b). (c) Number of DEGs between vegetative mycelium and other three developmental paths. (d) Venn diagram showing the DEGs shared by different developmental paths. (e) Volcano plots showing distributions of DEGs, taking gene expression levels in vegetative mycelium as reference, grey dots showing non-DEGs.

In this study, 10082 genes were detected to have expression in at least one developmental path, all of them can be matched with gene ID from strain Okayama-7 for further analysis (8, 9). Taking gene expression levels in vegetative mycelium as references, 3962 differentially expressed genes (DEGs) were detected in other three paths and assigned to 6 groups (3 paths with up-/down-regulation). Hyphal knot and sclerotia possessed larger amount of DEGs, with 1142 and 2065 up-regulated genes and 1011 and 1009 down-regulated genes, respectively (Fig. 3c). In oidia, only 175 genes were up-regulated, while 460 genes were down-regulated. 76 up-regulated and 171 down-regulated genes were shared by other three paths (Fig. 3d). Most DEGs displayed the fold change within 1/30 to 30 (Fig. 3e).

### Metabolic-related genes are differentially expressed in different developmental paths

To have a closer look on the transition of expression profiles in the developmental process, we performed Gene ortholog (GO) term and EuKaryotic Orthologous Groups (KOG) term enrich analysis according to the DEG groups. KOG enrichment profile showed a strong shift of gene set usage during the development of hyphal knot and sclerotia, while little regulations were performed during oidia formation (Fig. 4). Similar to the result of KOG enrichment, GO enrichment analysis present significant turn over on gene expression in hyphal knot (Fig. S2a). During hyphal knot formation, genes related to “biological processes” and “cellular component” were being down-regulated globally. Along the oidiation process, small number of DEGs were called and the functional enrichment was weak. These results show the high similarity on expression profiles of vegetative growth and oidiation. In sclerotia, down-regulated genes were enriched in molecular function group, indicating the lower metabolic rate at the sclerotia forming stage.

**Fig 4.**
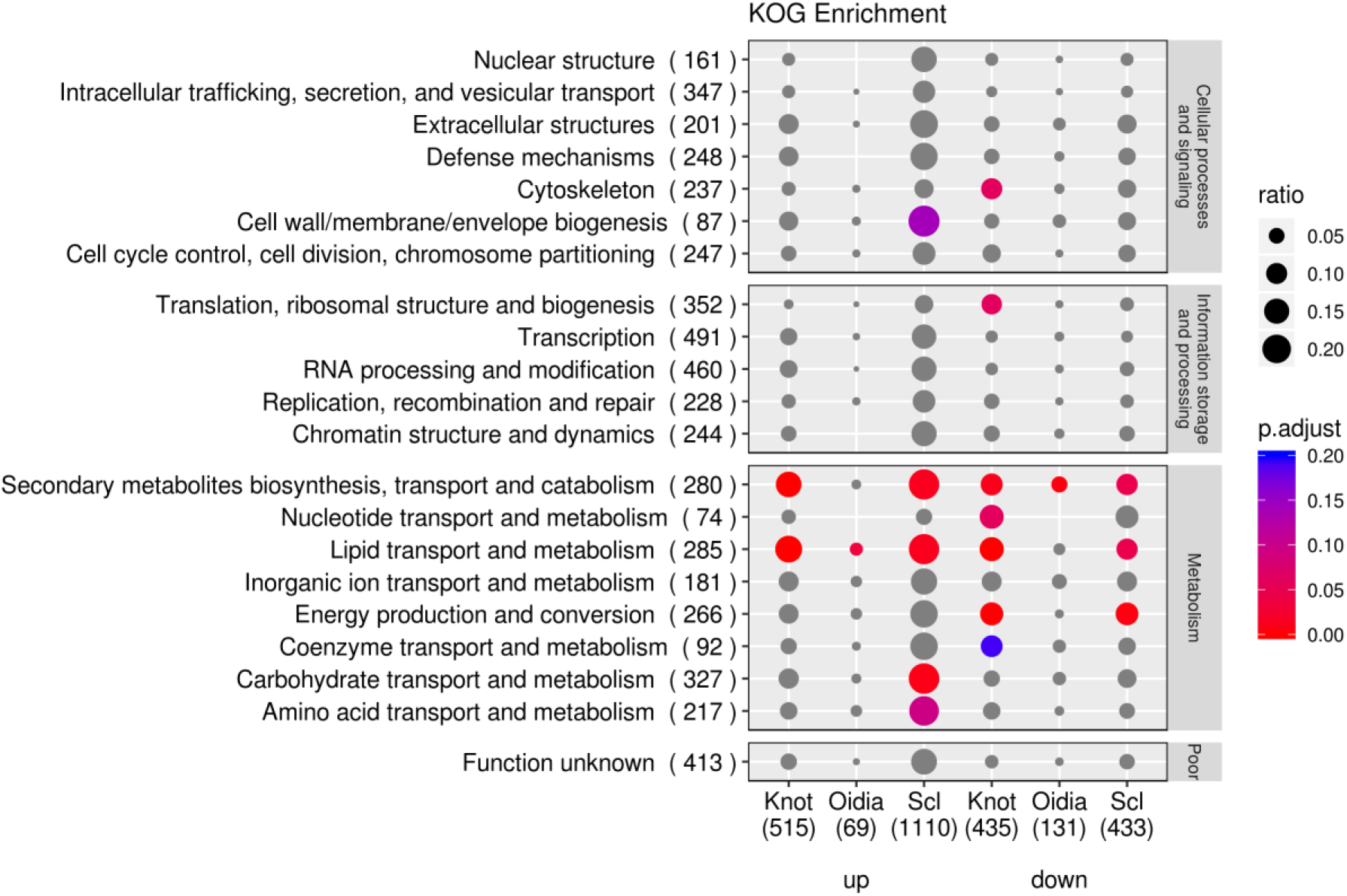
KOG enrichment analysis on DEGs on reproductive development. Number of annotated genes are listed below. Number beside KOG terms indicates number of genes being annotated to the node in the genome. Ratio is calculated by annotated genes of specific KOG term in each DEG group over annotated genes of specific KOG term in the genome background. Enrichment groups with Benjamini and Hochberg method (BH) adjusted p value ≤ 0.20 are shaded in red to blue color, others are in grey.

Functional analysis based on KOG terms reveals that down-regulated genes in both hyphal knot and sclerotia paths were enriched on energy production and conversion function. In sclerotia, genes with function of cell wall/membrane/envelope biogenesis, carbohydrate transport and metabolism and amino acid transport and metabolism were significantly up-regulated. GO term enrich analysis showed a significant enrichment on carbohydrate metabolic process and hydrolase activity of knot up-regulated, sclerotia up-regulated and oidia down-regulated genes. Expression level of “glucosidase” and “carboxy lyase activity” genes were up-regulated during hyphal knot and sclerotia development, while remained steady during oidia formation (Fig S3a). The shift on transcriptome indicates the re-distribution of materials and energy to fulfill the fruiting requirements. These results suggest that the determination of developmental paths is related to the internal regulation on carbohydrate metabolism and energy production and conversion.

### Gene expression in carbohydrate metabolic pathway explains carbon metabolic flux

Up-regulated genes in hyphal knot and sclerotia were significantly and strongly enriched on carbohydrate metabolic pathways (Fig. S2b). Here, we put the carbohydrate content assay results and gene expression results together (Fig. 5). Compare to vegetative mycelium, glycogen phosphorylase and glucoamylase were up-regulated in all three reproductive structure forming paths (Fig. S3b). These enzymes catalyze glycogen into glucose-1-P and free glucose, providing materials and energy for the formation of complex reproductive structures. The higher expression level of glycogen degrading enzymes, the lower glycogen content in the culture.

**Fig 5.**
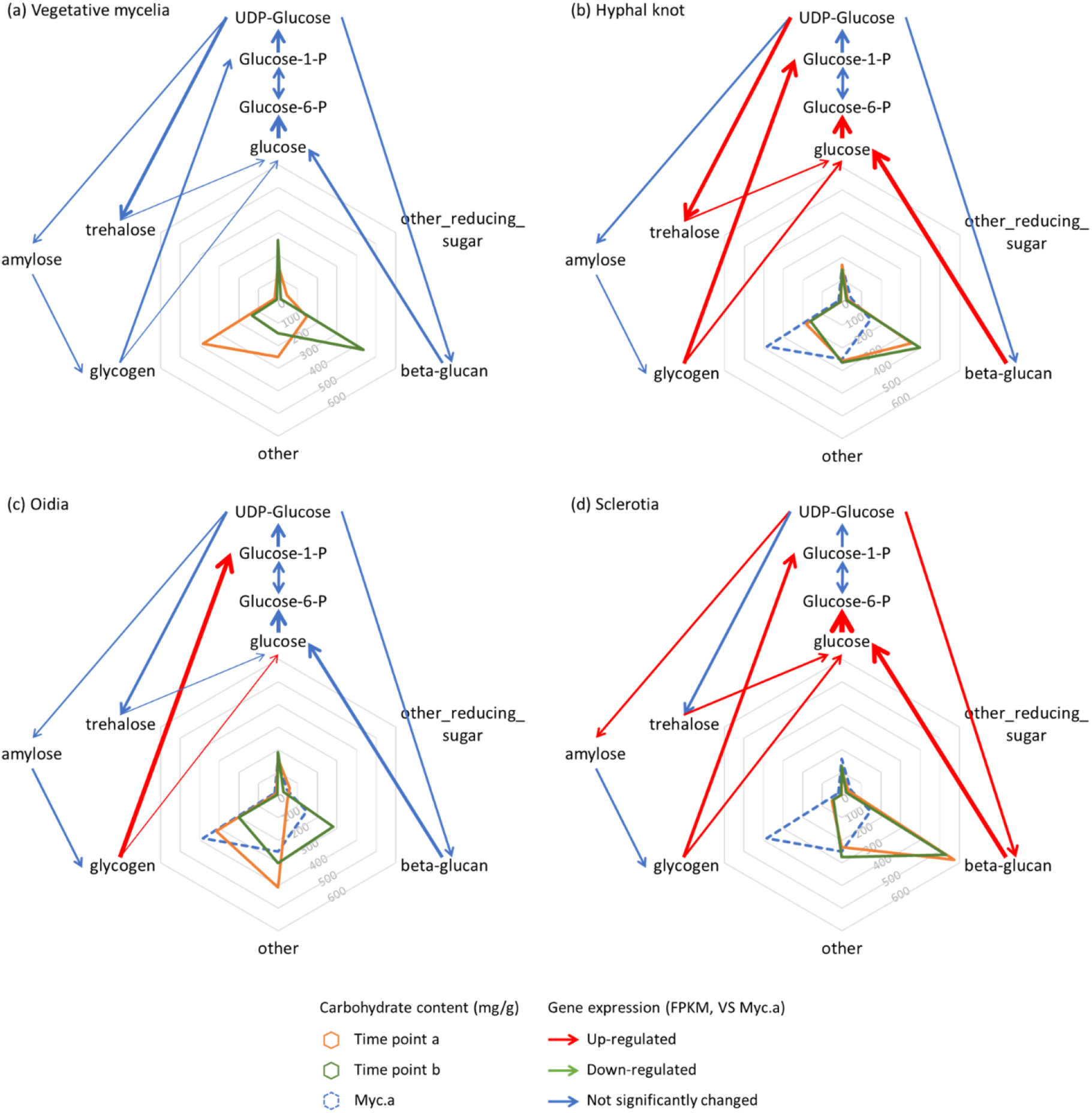
Gene expression level reflects the carbohydrate metabolic flux. Spider plots show content of different type of carbohydrate, arrows indicate the conversion from substrate to product. Thickness of the arrow indicates relative value of expression level of enzyme coding gene. Red arrows indicate up-regulation of expression level compare to vegetative mycelia, green arrows indicate down-regulation (none), and blue arrows indicate that the expression level is not significantly changed.

Unlike in the favorable environment that oidia are formed, during the fruiting process and sclerotia forming process, not only the glycogen catabolism, but also the metabolism of beta-glucan and trehalose increased. Higher relative expression level on beta-glucan synthase and beta-glucanase indicates the stronger carbon flow that run into beta-glucan anabolism during sclerotia formation. Such prediction coincided with the observation of stronger beta-glucan accumulation in sclerotia than hyphal knot. Although trehalose only took a tiny part of the dry weight in our study on *C. cinerea*, it is regarded as a carbon reserve and stress protectant in filamentous fungi, and found accumulating in resting cells such as spores and sclerotia (44, 45). Up-regulation on gene expression of trehalose phosphatase and trehalose phosphate synthase matched with the trehalose content in hyphal knot higher than sclerotia and oidia. Depletion of trehalose content in the latter developmental stages might be caused by the co-expression of trehalase. Up-regulation on both anabolism and catabolism of beta-glucan and trehalose suggest the high intensity of cell wall remodeling and complexity of multicellular structure formation (Fig. S4). In short, our sampling time point can represent the turning point of developmental differentiation. Conversion between glucose and glycogen and conversion between glucose and beta-glucan are the main carbon flows in the differentiation processes. The carbon metabolic flux can be explained on transcriptome level.

### Transcriptome age index and transcriptome divergence index profiles in developmental paths differentiation

To understand the transcriptome of developmental paths differentiation with evolutionary perspectives, we used transcriptome age index (TAI) and transcriptome divergence index (TDI) to estimate the evolutionary age and selective pressure of each developmental path (46–48). TAI and TDI was calculated based on either phylostrata (PS) or dN/dS ratio, and expression level of genes; the lower the TAI, the evolutionarily older the transcriptome; the lower the TDI with value less than 1, the stronger the force of purifying selection (48). The distributions of PS and dN/dS ratio of expressed genes in this study are shown in Fig. S6. We figured out that oidiation process expresses the evolutionarily oldest genes, next comes sclerotia, and vegetative mycelium and hyphal knot express the evolutionarily younger genes. All four developmental paths suffer from the strong selection force on the transcriptome. Oidia transcriptome emitted the strongest signal of purifying selection, and hyphal knot showed the weakest (Fig. 6a). Thus, oidia is the most conserved developmental path and fruiting process is evolutionarily young and divergent.

**Fig 6.**
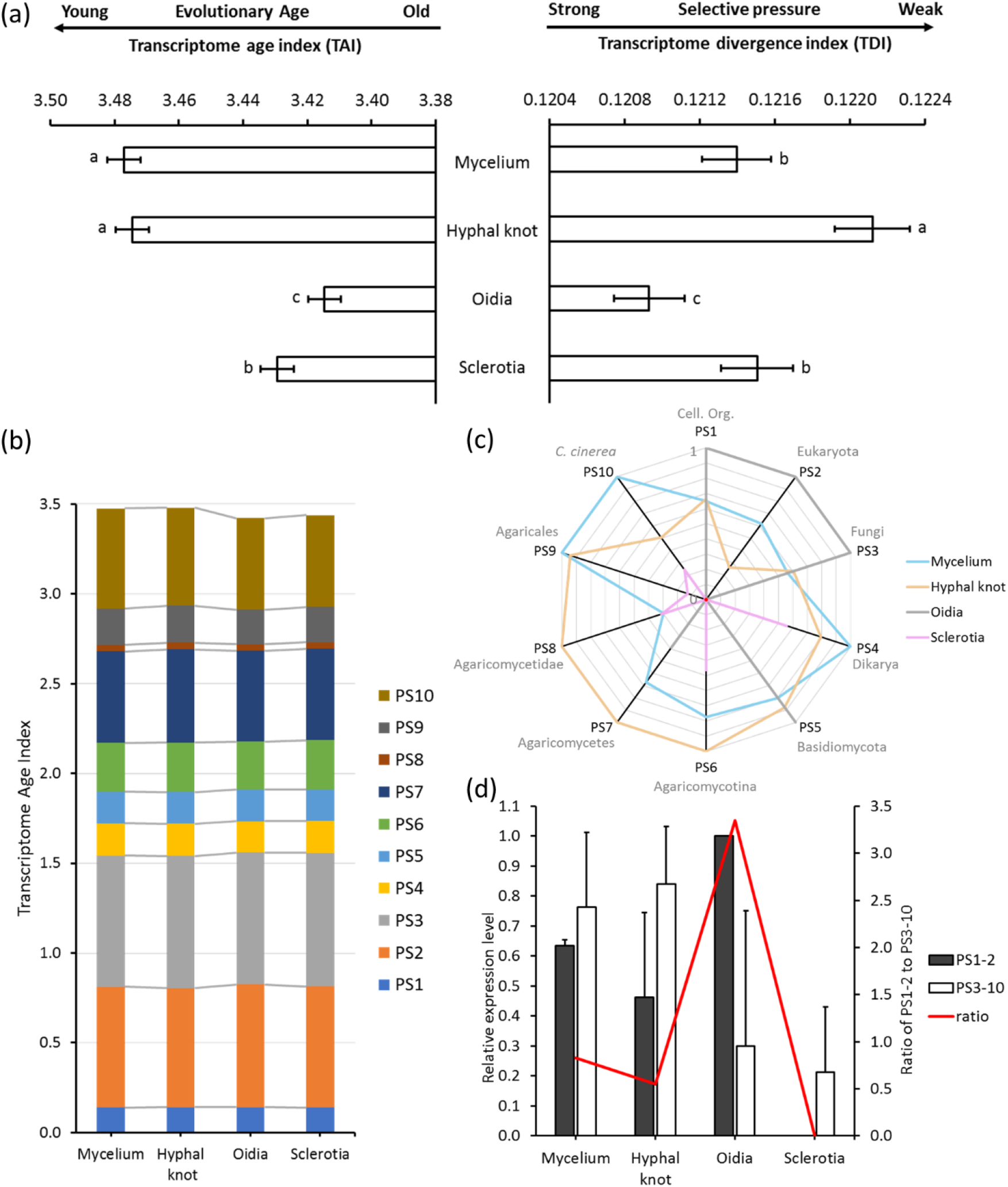
Evolutionary transcriptome profiles of developmental paths. (a) TAI and TDI values of different developmental paths. TAI quantifies the mean evolutionary age of a transcriptome. The lower the TAI, the evolutionarily older the transcriptome; TDI quantifies the mean selection force acting on a transcriptome. The lower the TDI, the stronger the force of purifying selection, giving its value less than 1. Error bars showing 95 % confidence interval estimated by bootstrap sampling for 1000 times. Lowercase letters showing TAI/TDI values that are significantly different among developmental paths in multiple comparisons (p < 0.05). (b) Contribution of each PS to the TAI: PS 3 > PS 2 > PS 10 > PS 7 > PS 6 > PS 9 > PS 4 ≈ PS 5 > PS 1 > PS 8. (c) Relative expression level of genes from each PS across developmental paths. (d) Mean relative expression level of old genes (PS 1-2) and young genes (PS 3-10) over developmental paths. Relative expression level (RE) of PS 1 and PS 2 in oidia are the same and equal to 1, RE of PS 1 and PS 2 in sclerotia are the same and equal to 0.

Among the ten phylostrata, PS 1-2 are defined as old genes and PS 3-10 are defined as young genes (48). Genes in PS 1-2 are in charge of basic cellular functions of eukaryotic cells. Intermediate phylostrata (PS 3-9) correspond to the divergence from fungi to Agaricales (gilled mushroom). Those genes mainly function on signal transduction and developmental regulation, and they involve in the extracellular structure formation, defense, transcription, RNA processing and modification, carbohydrate transportation and metabolism processes (Fig. S5).

Protein coding genes that first emerged in domain Eukaryota (PS 2), Fungi (PS 3) and *C. cinerea* (PS 10) contributed most to the TAI profile (Fig. 6b). Among four developmental paths, oidia had high expression level of old genes (PS 1 and PS 2) and young gene group PS 3 and PS 5, but low expression level of other younger genes of PS 6-10 (Fig. 6c). On the contrary, hyphal knot displayed high expression level on PS 5-9, the gilled mushroom-specific genes, but low expression level in PS 1-3. Vegetative mycelium had high expression level of PS 9-10. Genes from these two phylostrata are Agaricales-specific or *C. cinerea* unique genes. These young genes display the functional enrichment on defense and carbohydrate binding. In sclerotia, most of the metabolic processes slowed down under stress. Sclerotia got the lowest expression level of old genes and relatively lower expression level of young genes. The highest expression ratio of old genes to young genes occurred in oidia, followed by mycelium and hyphal knot, and sclerotia had the lowest ratio of zero (Fig. 6d). Interestingly, such ranking coincided with the evaluation on growing environment, from the most temperate to the most stressful. All these results indicate that oidiation, the asexual reproductive process, is generated from the common ancestor of eukaryota and conserved in current living organisms; while fruiting, the sexual reproductive process, is highly specific and evolutionarily adaptive.

### RNA editing events happened in all developmental paths

In this study, a total of 245 RNA editing sites and 819 RNA editing events were identified in 4 developmental paths. The RNA editome in developmental paths differentiation of *C. cinerea* is strongly different from those Ascomycetes fungi. The majority of RNA editing events had editing levels < 20 % (Fig. 7a), and the editing level remained constant and relatively low (Fig. 7b). T-to-C substitution took up over 55 % of the editing events (Fig. 7c). 107, 111, 101 and 171 editing sites were detected in mycelium, hyphal knot, oidia and sclerotia, respectively (Fig. 7d). In *C. cinerea*, 44.1 % the RNA editing sites were annotated to the intergenic region and only 30.4 % are in the coding sequence (CDS, Fig. 7e). RNA editing events tend to appear on genes that are evolutionarily old and under strong purifying selection (Fig. S6).

**Fig 7.**
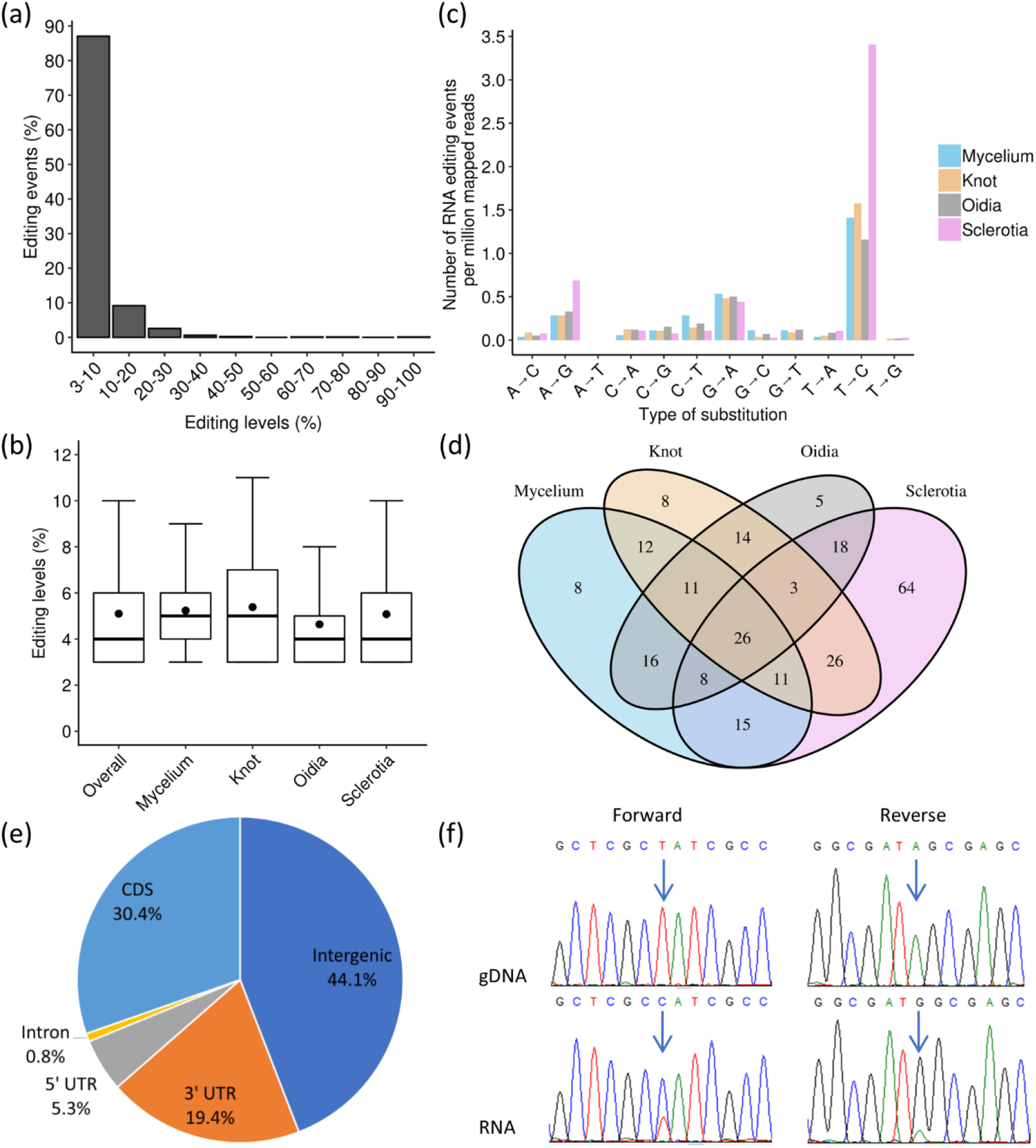
Properties of RNA editing sites and RNA editing events. (a) Histogram showing editing the frequency of 819 RNA editing events. (b) The number of each type of RNA editing events per million mapped reads in different developmental stages. (c) Box plots showing RNA editing levels of RNA editing events in different developmental paths. (d) Venn diagram showing the number of RNA editing sites shared by different developmental paths. (e) The distribution of 245 RNA editing sites. (f) Sanger sequencing validates T-to-C RNA editing events on scaffold_131:54938 (- strand), blue arrow indicates editing site.

To validate the RNA editing events, a selection of candidate sites was chosen to be amplified and sequenced using Sanger method. Site scaffold_131:54938 was found edited in 9 of 12 biological samples, with editing level of 3 – 70 %. Clear signal of double peaks showing T-to-C editing can be observed in 6 samples and the intensity of signals were consistent to the RNA-seq results (Fig. 7f and Table S2). Single sequencing signal peak was observed in DNA and predicted unedited samples. This editing site is on a hypothetical protein (CC1G_15451), and it is a synonymous variant on the transcript that codes alanine.

### Potential impact of RNA editing on gene expression regulation

RNA editing site-containing genes were grouped according to their editing impact annotation and performed KOG enrichment analysis (Fig. S7). 31 and 40 RNA editing sites in CDS can cause either synonymous or nonsynonymous changes of the protein sequence. Six editing sites can generate splicing variants and transcript length variants (Table 1). Four of these editing containing genes, including anthranilate phosphoribosyltransferase, manganese superoxide dismutase, glyoxalase I and DNA-directed RNA polymerase II, only have single copy in the genome and all with unique function.

**Table 1.** RNA editing events result in splicing variants and transcript length variants.

| **RNA editing position** | **Editing impact** | **Annotation** | **Okayama-7 #130 ID** |
| --- | --- | --- | --- |
| scaffold_250:38334 | Splice acceptor variant and intron variant | Anthranilate phosphoribosyltransferase | CC1G_02982 |
| scaffold_194:26812 | Splice region variant and intron variant | DNA-directed RNA polymerase II | CC1G_02497 |
| scaffold_42:66264 | Splice region variant | Manganese superoxide dismutase | CC1G_03559 |
| scaffold_124:43286 | Splice region variant | Glyoxalase I | CC1G_02307 |
| scaffold_239:36981 | Splice region variant | Hypothetical protein | CC1G_12295 |
| scaffold_78:81092 | Stop gained | Trp-Asp repeats containing protein | CC1G_03748 |

MicroRNAs could mediate post-transcriptional gene expression regulation by base pairing their seed region (2-7 nt at the 5’-end) to the UTR (49). RNA editing in 3’ UTR can create or destroy the microRNA recognition sites, resulting in the changes of mRNA degradation and translational repression (50). 35 micro-RNA like RNA (milRNA) had been predicted in *C. cinerea* strain #326 and 7 of them were validated by reverse transcription-qPCR (Lau *et al*, unpublished data). Among 48 editing sites that locate in 3’ UTR, 18 of them were predicted to interact with the known milRNA (Table 2). The T-to-C editing event at 3’ UTR of thiamine biosynthetic bifunctional enzyme (scaffold_76:107237, CC1G_03317) could cause binding loss of validated milRNA cci-milR-32-3p and cci-milR-33-3p, and binding gain of predicted milRNA cci-milR-32-5p. Another T-to-C editing event at 3’ UTR of glyceraldehyde-3-phosphate dehydrogenase (scaffold_77:35473, CC1G_09116) would create the milRNA recognition site for the validated milRNA cci-milR-22.

**Table 2.** Interaction between RNA editing sites and milRNA. * for RT-qPCR validated milRNA.

| RNA editing<br>position | miRNA binding |  |  |
| --- | --- | --- | --- |
|  | Lost | Gain | Remains |
| scaffold_105:15293 |  | cci-miR-38-5p |  |
| scaffold_118:64567 | cci-miR-33-3p* |  | cci-miR-27 |
| scaffold_118:64602 | cci-miR-21a-5p,<br>cci-miR-21b-5p |  |  |
| scaffold_13:185033 | cci-miR-28 | cci-miR-33-5p |  |
| scaffold_23:18866 |  |  | cci-miR-18,<br>cci-miR-33-3p* |
| scaffold_23:198001 |  |  | cci-miR-25,<br>cci-miR-32-3p*,<br>cci-miR-33-3p* |
| scaffold_3:265899 |  |  | cci-miR-32-3p* |
| scaffold_302:10673 | cci-miR-33-5p,<br>cci-miR-35 |  |  |
| scaffold_320:19744 |  |  | cci-miR-25 |
| scaffold_340:18068 | cci-miR-33-5p |  |  |
| scaffold_343:9606 |  |  | cci-miR-17* |
| scaffold_390:3277 |  |  | cci-miR-33-3p*,<br>cci-miR-33-5p |
| scaffold_412:12417 |  |  | cci-miR-30 |
| scaffold_71:107334 |  |  | cci-miR-19-3p |
| scaffold_73:108325 | cci-miR-23 |  |  |
| scaffold_76:107237 | cci-miR-32-3p*,<br>cci-miR-33-3p* | cci-miR-32-5p |  |
| scaffold_76:45473/6 |  | cci-miR-27 |  |
| scaffold_77:35473 |  | cci-miR-22* |  |

## Discussion

As a model mushroom-forming fungus, fruiting process of *C. cinerea* is well studied physiologically and genetically. In this study, we monitor the early developmental process of three destinies, fruiting, oidiation and sclerotia formation, with the reference of vegetative mycelium. Carbohydrate assays and high-throughput sequencing results all indicate the essential role of carbon metabolic flux in fungal morphogenesis. RNA editing, a co-/post-transcriptional modification which could rewrite the information in RNA, also prefers carbohydrate transport and metabolism transcripts. Evolutionary transcriptome analysis reveals the origin and selection of dispersal and survival strategies of *C. cinerea* (Fig. 8).

**Fig 8.**
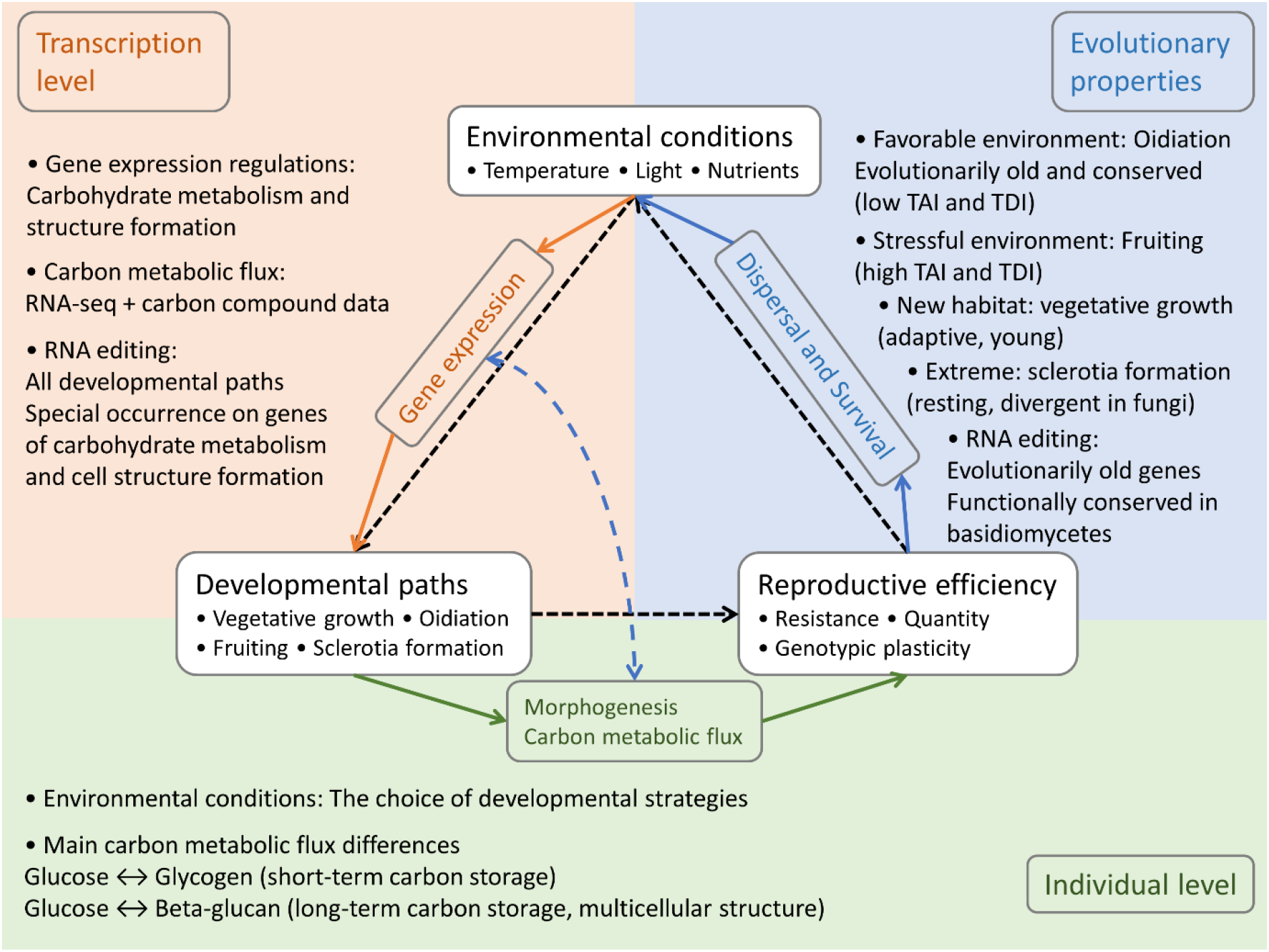
Properties of developmental paths differentiation in *C. cinerea*

Fruiting plays a pivotal role in fungal life cycle and attracts research attention (17, 51, 52). But we also need to know the role of oidiation and sclerotia formation in optimizing the dispersal and survival fitness of fungi under divergent environmental conditions (53). In this study, the most optimal to the most stressful environment for fungal development are ranked as the incubation conditions of oidiation, vegetative growth, fruiting and sclerotia formation. Nutrient, temperature and light are three critical environmental factors for the fruiting body development in mushroom-forming fungi (51). Light indicates open space to the fungi. In ascomycetes, light inhibits sexual reproduction but induces asexual development and produce tremendous amount of conidia (54). In *C. cinerea*, the combination of rich nutrient, optimum temperature and illumination results in the high production of genetically identical asexual spores, the oidia (40). Oidia contain ready-to-use materials for the rapid gemination, and with little resistant protective structures (55). Its high sensitivity to environmental stress will restrict the dispersal of fungi (56). Oidiation process is energy-efficient but has little genetic flexibility. Oidia disperse rapidly in rare environments. The lack of light indicates the lack of open air (57). Under such circumstances, to grow connected hyphae, instead of releasable oidia, would be a more promising way to expand the habitat. When nutrient depletes, fungi changes their developmental strategies for long-distance dispersal and long-term survival (2, 16, 58). Fruiting and sclerotia formation are both energy consuming processes because of the difficulty in forming highly organized multicellular structure (59). Thus, environmental stress like nutrient depletion is essential for the induction of fruiting and sclerotia formation. Basidiospores and sclerotia possess thicker resistant layer than oidia and hyphae, and both of them are able to travel in distance and last for years (56, 60). Comparing to glycogen, beta-glucan is much difficult to be used or digested by other organisms. Storing beta-glucan as the carbon source for future usage is an energy-efficient choice in basidiospores and sclerotia (61–63). Like oidia, sclerotia also have little genetic flexibility. On the contrary, basidiospores benefit from the sexual propagation and high genotypic plasticity (64, 65). Thus, basidiospores would adapt to the new habitat more successfully.

Evolutionary transcriptomic analysis has been performed in animal and plant embryogenesis and fungal fruiting (48, 66). The phylotranscriptomic hourglass pattern across kingdom illustrates the occurrence of complex multicellularity. Our results uncovered the correlation of environmental stress, developmental destinies and gene evolution in the life history of *C. cinerea*. Oidiation appears in highly favorable environment, genes express in this process are mainly those common to all eukaryotes and they are under strong purifying selection. Regulation on gene expression ensure the unspecified process of oidia formation to be performed accurately and sufficiently. Vegetative mycelium grows in changeable environment and it is in charge of primary occupation of new habitat. The Agaricales-specific and species unique genes are very likely to benefit the fungus on niche segmentation and adapting to the complex and changing environment during vegetative growth (38, 67, 68). High expression on species-specific genes and relative lower selective pressure of purifying selection enable mycelium to minimize the interspecific competition and improve fitness (69). Hyphal knots display similar transcriptome age as vegetative mycelium, but the former has higher transcriptome divergence. These indicate that the developmental process of fruiting body is relatively conserved, but genes of specific biological functions got diversity across genus (17, 48). Sclerotia function as the persistent resting structure which would need to suffer the extreme stressful environment (58). The relative old transcriptome age and strong purifying selection force of sclerotia suggest the common demand of overcoming stress and the importance of keeping such function in the genome for fungal species (70–72). The expression profile with phylostrata also indicates that the sclerotia formation process is highly divergent and with little similarity among different fungi (58, 70, 72).

As a special kind of transcriptional modification, RNA editing has been widely found in all domains of life and showed significant impacts on diverging the transcriptome and proteome (73–76). In fungi, developmentally regulated for sexual reproduction and evolutionarily conserved across genus are two main patterns of RNA editing events (35, 37, 38). Our results show that RNA editing not only occurs in fruiting, but also other developmental paths. Similar to the stage-specific and substrate-specific RNA editing studies on basidiomycetes, editing events in *C. cinerea* regulate the genes in carbohydrate metabolism and cell structure formation (37, 38). All these results emphasize the functional similarity of RNA editing in regulating carbohydrate metabolism, and further, the essential role of carbon metabolism in fungal development. RNA editing show preference on genes that are old and under high purifying selection, revealing the special roles of RNA editing on providing plasticity to the transcriptome (34).

The profile of RNA editome of *C. cinerea* is similar to the previous studies in basidiomycetes, showing weak A-to-I editing preference (37, 38, 77). However, the number of called RNA editing events is significantly less. Moreover, compare to a basidiomycetes *G. lucidum* and two ascomycetes, *F. graminearum* and *N. crassa*, that over 60 % of the RNA editing sites are predicted to be in the CDS (35, 37, 78), only around 30 % of the RNA editing sites are in CDS in *C. cinerea*. Despite the differences of species from distant order, we refer that the low editing density is caused by the usage of a homokaryotic strain, also by the much stricter filtering parameters. In addition, we resequenced the genome of strain #326 as a reference to exclude false positive events introduced by inaccuracy in genome assembly and genome variants. We have blast out 3 adenosine deaminases acting on tRNA (ADATs) homologs in the genome, but no adenosine deaminases acting on RNA (ADARs) homolog (37, 78). The mechanism of RNA editing in *C. cinerea* remains unknown.

## Conclusions

Responding to the environmental conditions, *C. cinerea* follows different developmental paths including vegetative growth, oidiation, fruiting and sclerotia formation. In these paths, genes are differentially expressed and RNA editing occurred. Carbohydrate metabolism is strictly regulated and differs dramatically. Glycogen is produced as storage carbon source to provide energy in the early developmental stages. Stored glycogen is converted into beta-glucan during fruiting or sclerotia formation. Carbon metabolic flux is regulated to fulfill the demands of short-term usage and long-term survival adapting to the specific environmental conditions. Phylotranscriptomic analysis showed that oidiation happens in favorable environments, and has the oldest transcriptome age and the lowest transcriptome divergence. Developmental paths of fruiting and sclerotia formation that occur in stress environments have evolutionarily younger genes expressed, and display younger transcriptome age and higher transcriptome divergence. Editome analysis showed that RNA editing occurs in all developmental paths and is developmentally regulated. RNA editing appears to regulate carbohydrate metabolism genes at both transcriptional and post-transcriptional levels. These events provide more plasticity to the transcriptome and show preferences on conserved genes. In short, the differentiation of developmental paths in *C. cinerea* is regulated transcriptionally and post-transcriptionally, with major changes in carbon metabolic flux.

## Materials and methods

### Strain

The homokaryotic fruiting strain *C. cinerea* #326 (A43mut B43mut pab1-1) was grown at 37 °C on solid YMG medium for 5 days before reaching the edge of the 90 mm petri dish to obtain the working stock plates. Culture condition, growth rate measurement, and sample collection for carbohydrate content determination and RNA sequencing are described in supplementary methods (Text S1).

### RNA-seq library preparation and sequencing

RNA with three biological replicates of each path were proceeded to RNA-seq. About 5 µg of total RNA for each sample was sent to the Beijing Genomics Institute (BGI, Shenzhen, China) for library construction and sequencing. The unstranded RNA library was prepared using TruSeq RNA Sample Prep Kit v2 (Illumina, USA), and sequenced with Illumina HiSeq® 4000 at the 2×150 bp paired-end read mode. *In silico* analysis on transcriptome, including reads alignment, DEG detection, functional classification and phylotranscriptomic analysis, is described in supplementary methods (Text S1). RNA editing sites was called by REDItools v1.0.4 (79), parameters used are also listed in supplementary methods (Text S1).

### DNA extraction and genome resequencing

Genomic DNA was extracted from vegetative mycelium of *C. cinerea* using DNeasy Plant Mini Kit (Qiagen, Germany). DNA sequencing library was prepared with insert size of 270 bp and sequenced with Illumina HiSeq® 4000 at the 2×150 bp paired-end read mode. 9.2 million clean reads and 1.38 billion clean bases were obtained. Reads were aligned to the reference genome of *C. cinerea* strain #326 released in Genome portal of Joint Genome Institute (https://mycocosm.jgi.doe.gov/Copci_AmutBmut1/Copci_AmutBmut1.home.html) using Bowtie2 (80).

### qRT-PCR validation

Quantitative real-time PCR (qRT-PCR) was used to validate the RNA-seq results. cDNA was synthesized from total RNA with anchored-oligo(dT)18 primer and random hexamer primer using Transcriptor First Strand cDNA Synthesis Kit (Roche, Germany). The template-primer mix was denaturized at 65 °C, and the RT reaction was incubated as follows: 10 min at 25 °C, 30 min at 55 °C and 5 min at 85 °C. 1 µg of RNA was input into each 20 µl RT reaction. Several genes were selected, and the expression was quantitatively measured using SsoAdvanced^TM^ Universal SYBR® Green Supermix (Bio-Rad, USA) with Applied Biosystems^TM^ 7500 fast Real-Time PCR System (Applied Biosystems, USA). PCR reactions were performed as the following program: 30 sec at 95 °C, followed by 40 cycles 15 sec at 95 °C and 30 sec at 60 °C, instrument default setting on melt-curve analysis. 18S is used as the internal control. Primer used in this study are listed in supplementary methods (Text S1).

### Validation of RNA editing sites

PCR amplification of RNA editing containing sequence were performed by using cDNA and gDNA as templates with KAPA HiFi HotStart ReadyMix PCR kit (Roche, Germany) and the following program: 95 °C for 3 min, followed by 30 cycles 98 °C for 20 sec, 65 °C for 20 sec, and 72 °C for 15 sec, and 72 °C for 1 min. PCR products are detected on 1.5 % agarose gel and purified with MEGA quick-spin Plus Fragment DNA Purification Kit (MEGA, Korea). Sanger sequencing of PCR products were performed by BGI.

## Data availability

Sequencing data of this study have been submitted to the NCBI Sequence Read Archive (SRA, http://www.ncbi.nlm.nih.gov/sra) under BioProject accession numbers PRJNA573619 and PRJNA573620.

## Acknowledgements

We would like to thank Dr. Hajime Muraguchi for kindly sharing the *C. cinerea* strain #326 with us. We would also like to thank Xuanjin Cheng and Man Kit Cheung for the fruitful discussion.

This work was supported by the RGC General Research Fund (GRF 14116916 and GRF 14103817) provided by the Research Grants Council of HKSAR, PRC. The funders had no role in study design, data collection and interpretation, or the decision to submit the work for publication.

## Supplementary materials Supplementary text

Text S1. Supplementary methods.

### Supplementary tables

Table S1. Carbohydrate content in different developmental paths. Multiple comparison results are shown with groups.

Table S2. Validation of RNA editing events with Sanger sequencing. RNA editing candidate scaffold_131:54938 (- strand), T-to-C editing show in red box. * for RNA editing events being significantly observed.

### Supplementary figures

Fig S1.Growth status of colonies. oidia production (a) and mycelium growth (b) when incubated in 37 °C with continuous light or continuous dark.

Fig S2. GO (a) and KEGG (b) enrichment analysis on DEGs on reproductive development. Number of annotated genes are listed below each DEG group. Number beside GO terms indicates number of genes being annotated to the node in the genome. Ratio is calculated by annotated genes of specific GO term in each DEG group over annotated genes of specific GO term in the genome background. Enrichment groups with Benjamini and Hochberg method (BH) adjusted p value ≤ 0.20 are shaded in red to blue color, others are in grey.

Fig. S3. (a) Log2FPKM of genes on starch and sucrose metabolism pathway cci00500. (b) qPCR validation of gene expression on carbohydrate metabolic pathway related genes.

Fig S4. Log2 fold change values of genes related to KOG group of (a) carbohydrate transport and metabolism; (b) energy production and conversion; (c) mycelial structure formation.

Fig. S5. KOG enrichment analysis (a) and GO enrichment analysis (GO term level 3, b) of phylostrata. Number below each PS shows number of annotated genes in specific PS. Number beside KOG/GO terms indicates number of genes being annotated to the node in the genome. Ratio is calculated by genes of specific PS annotated to the KOG/GO term over annotated genes of specific PS. For GO enrichment, ratio ≥ 0.1 are plotted. Enrichment groups with Benjamini and Hochberg method (BH) adjusted p value ≤ 0.20 are shaded in red to blue color, others are in grey.

Fig S6. Evolutionary transcriptome and RNA editome profiles of C. cinerea development. Distribution of Phylostrata (a) and dN/dS ratio (b) of all expressed genes (upper panel) and RNA editing site containing genes (lower panel). Red dot line showing the mean. RNA editing site containing genes are evolutionarily older in the genome. (c) and (d) shows percentage of RNA editing containing genes in each phylostrata or dN/dS group.

Fig S7. KOG enrichment analysis of RNA editing sites. 245 editing sites in total. Number before each annotation category shows number of RNA editing sites with specific functional annotation predicted by snpEff. Number beside KOG terms indicates number of genes being annotated to the node in the genome. Ratio is calculated by genes of specific functional category annotated to the KOG term over annotated genes of specific KOG term in the genome background. Enrichment groups with Benjamini and Hochberg method (BH) adjusted p value ≤ 0.20 are shaded in red to blue color, others are in grey.

## References

1. Raudaskoski M. 2015. Mating-type genes and hyphal fusions in filamentous basidiomycetes. Fungal Biol Rev 29:179–193.

2. Halbwachs H, Simmel J, Bässler C. 2016. Tales and mysteries of fungal fruiting: How morphological and physiological traits affect a pileate lifestyle. Fungal Biol Rev 30:36– 61.

3. Sipos G, Prasanna AN, Walther MC, O’Connor E, Balint B, Krizsan K, Kiss B, Hess J, Slot J, Riley R, Boka B, Rigling D, Barry K, Lee J, Mihaltseva S, Labutti K, Lipzen A, Waldron R, Moloney N, Sperisen C, Kredics L, Vagvolgyi C, Patrigniani A, Fitzpatrick D, Nagy I, Doyle S, Anderson JB, Grigoriev I V., Guldener U, Munsterkotter M, Varga T, Nagy LG. 2017. Genome expansion and lineage-specific genetic innovations in the world’s largest organisms (Armillaria). Nat Ecol Evol 1:1931–1941.

4. Kües U. 2000. Life history and developmental processes in the basidiomycete *Coprinus cinereus*. Microbiol Mol Biol Rev 64:316–53.

5. Gomez-Zavaglia A, Shaohua S, Zeng F, Zhu W, Zhang S, Hu B, Wei W, Xiong Y, Peng F, Yu Y, Zheng Y, Chen P. 2016. *De novo* analysis of *Wolfiporia cocos* transcriptome to reveal the differentially expressed carbohydrate-active enzymes (CAZymes) genes during the early stage of sclerotial growth. Front Microbiol 7:83.

6. Rodenburg SYA, Terhem RB, Veloso J, Stassen JHM, van Kan JAL. 2018. Functional analysis of mating type genes and transcriptome analysis during fruiting body development of *Botrytis cinerea*. MBio 9:e01939–17.

7. Erental A, Dickman MB, Yarden O. 2008. Sclerotial development in *Sclerotinia sclerotiorum*: awakening molecular analysis of a “Dormant” structure. Fungal Biol Rev 22:6–16.

8. Stajich JE, Wilke SK, Ahrén D, Au CH, Birren BW, Borodovsky M, Burns C, Canbäck B, Casselton LA, Cheng CK, Deng J, Dietrich FS, Fargo DC, Farman ML, Gathman AC, Goldberg J, Guigó R, Hoegger PJ, Hooker JB, Huggins A, James TY, Kamada T, Kilaru S, Kodira C, Kües U, Kupfer D, Kwan HS, Lomsadze A, Li W, Lilly WW, Ma L-J, Mackey AJ, Manning G, Martin F, Muraguchi H, Natvig DO, Palmerini H, Ramesh MA, Rehmeyer CJ, Roe BA, Shenoy N, Stanke M, Ter-Hovhannisyan V, Tunlid A, Velagapudi R, Vision TJ, Zeng Q, Zolan ME, Pukkila PJ. 2010. Insights into evolution of multicellular fungi from the assembled chromosomes of the mushroom *Coprinopsis cinerea* (*Coprinus cinereus*). Proc Natl Acad Sci U S A 107:11889–94.

9. Muraguchi H, Umezawa K, Niikura M, Yoshida M, Kozaki T, Ishii K, Sakai K, Shimizu M, Nakahori K, Sakamoto Y, Choi C, Ngan CY, Lindquist E, Lipzen A, Tritt A, Haridas S, Barry K, Grigoriev I V., Pukkila PJ. 2015. Strand-specific RNA-seq analyses of fruiting body development in *Coprinopsis cinerea*. PLoS One 10:e0141586.

10. Nunes LR, Costa de Oliveira R, Leite DB, da Silva VS, dos Reis Marques E, da Silva Ferreira ME, Ribeiro DCD, de Souza Bernardes LA, Goldman MHS, Puccia R, Travassos LR, Batista WL, Nóbrega MP, Nobrega FG, Yang D-Y, de Bragança Pereira CA, Goldman GH. 2005. Transcriptome analysis of *Paracoccidioides brasiliensis* cells undergoing mycelium-to-yeast transition. Eukaryot Cell 4:2115–28.

11. Fu Y, Dai Y, Yang C, Wei P, Song B, Yang Y, Sun L, Zhang Z-W, Li Y. 2017. Comparative transcriptome analysis identified candidate genes related to Bailinggu mushroom formation and genetic markers for genetic analyses and breeding. Sci Rep 7:9266.

12. Wen J, Zhang Z, Gong L, Xun H, Li J, Qi B, Wang Q, Li X, Li Y, Liu B. 2019. Transcriptome changes during major developmental transitions accompanied with little alteration of DNA methylome in two *Pleurotus* species. Genes (Basel) 10:465.

13. Dörnte B, Kües U. 2012. Reliability in transformation of the basidiomycete *Coprinopsis cinerea*. Curr Trends Biotechnol Pharm 6:340–355.

14. Srivilai P, Loutchanwo P. 2009. *Coprinopsis cinerea* as a model fungus to evaluate genes underlying sexual development in basidiomycetes. Pakistan J Biol Sci 12:821–835.

15. Watanabe T, Tagawa M, Tamaki H, Hanada S, Watanabe T, Tagawa ÁM, Tamaki ÁH, Hanada ÁS. 2011. *Coprinopsis cinerea* from rice husks forming sclerotia in agar culture. Mycoscience 52:152–156.

16. Moore D, Jirjis RI. 1976. Regulation of sclerotium production by primary metabolites in *Coprinus cinereus* (= *C. lagopus* sensu lewis). Trans Br Mycol Soc 66:377–382.

17. Krizsán K, Almási É, Merényi Z, Sahu N, Virágh M, Kószó T, Mondo S, Kiss B, Bálint B, Kües U, Barry K, Cseklye J, Hegedüs B, Henrissat B, Johnson J, Lipzen A, Ohm RA, Nagy I, Pangilinan J, Yan J, Xiong Y, Grigoriev I V, Hibbett DS, Nagy LG. 2019. Transcriptomic atlas of mushroom development reveals conserved genes behind complex multicellularity in fungi. Proc Natl Acad Sci U S A 116:7409–7418.

18. Daran-Lapujade P, Jansen MLA, Daran J-M, van Gulik W, de Winde JH, Pronk JT. 2004. Role of transcriptional regulation in controlling fluxes in central carbon metabolism of *Saccharomyces cerevisiae*. A chemostat culture study. J Biol Chem 279:9125–38.

19. Carreira R, Evangelista P, Maia P, Vilaça P, Pont M, Tomb J-F, Rocha I, Rocha M. 2014. CBFA: phenotype prediction integrating metabolic models with constraints derived from experimental data. BMC Syst Biol 8:123.

20. Lagziel S, Lee WD, Shlomi T. 2019. Studying metabolic flux adaptations in cancer through integrated experimental-computational approaches. BMC Biol 17:51.

21. Traven A, Jänicke A, Harrison P, Swaminathan A, Seemann T, Beilharz TH. 2012. Transcriptional profiling of a yeast colony provides new insight into the heterogeneity of multicellular fungal communities. PLoS One 7:e46243.

22. Busch S, Braus GH. 2007. How to build a fungal fruit body: from uniform cells to specialized tissue. Mol Microbiol 64:873–876.

23. Andoh T, Hirata Y, Kikuchi A. 2000. Yeast Glycogen Synthase Kinase 3 Is Involved in Protein Degradation in Cooperation with Bul1, Bul2, and Rsp5. Mol Cell Biol 20:6712– 6720.

24. Bidochka MJ, Low NH, Khachatourians GG. 1990. Carbohydrate storage in the entomopathogenic fungus *Beauveria bassiana*. Appl Environ Microbiol 56:3186–90.

25. Brunt IC, Moore D. 1989. Intracellular glycogen stimulates fruiting in *Coprinus cinereus*. Mycol Res 93:543–546.

26. Jirjis RI, Moore D. 1976. Involvement of glycogen in morphogenesis of *Coprinus cinereus*. J Gen Microbiol 95:348–352.

27. Qin J, Wang G, Jiang C, Xu J-R, Wang C. 2015. Fgk3 glycogen synthase kinase is important for development, pathogenesis, and stress responses in *Fusarium graminearum*. Sci Rep 5:8504.

28. Chen S, Xu J, Liu C, Zhu Y, Nelson DR, Zhou S, Li C, Wang L, Guo X, Sun Y, Luo H, Li Y, Song J, Henrissat B, Levasseur A, Qian J, Li J, Luo X, Shi L, He L, Xiang L, Xu X, Niu Y, Li Q, Han M V, Yan H, Zhang J, Chen H, Lv A, Wang Z, Liu M, Schwartz DC, Sun C. 2012. Genome sequence of the model medicinal mushroom *Ganoderma lucidum*. Nat Commun 3:913.

29. Sakamoto Y, Nakade K, Konno N. 2011. Endo-β-1,3-glucanase GLU1, from the fruiting body of *Lentinula edodes*, belongs to a new glycoside hydrolase family. Appl Environ Microbiol 77:8350–4.

30. Fukuda K, Hiraga M, Asakuma S, Arai I, Seikikawa M, Urashima T. 2008. Purification and characterization of a novel Exo-β-1,3-1,6-glucanase from the fruiting body of the edible mushroom Enoki (*Flammulina velutipes*). Biosci Biotechnol Biochem 72:3107– 3113.

31. Wang J, Zhang W, Zhou Y, Yuan S, Liu Z. 2015. Purification, characterization and synergism in autolysis of a group of 1,3-β-glucan hydrolases from the pilei of *Coprinopsis cinerea* fruiting bodies. Microbiology 161:1978–1989.

32. Nowrousian M. 2018. Genomics and transcriptomics to study fruiting body development: An update. Fungal Biol Rev 32:231–235.

33. Horton TL, Landweber LF. 2002. Rewriting the information in DNA: RNA editing in kinetoplastids and myxomycetes. Curr Opin Microbiol 5:620–6.

34. Liscovitch-Brauer N, Alon S, Porath HT, Elstein B, Unger R, Ziv T, Admon A, Levanon EY, Rosenthal JJC, Eisenberg E. 2017. Trade-off between transcriptome plasticity and genome evolution in cephalopods. Cell 169:191–202.e11.

35. Liu H, Li Y, Chen D, Qi Z, Wang Q, Wang J, Jiang C, Xu J-R. 2017. A-to-I RNA editing is developmentally regulated and generally adaptive for sexual reproduction in *Neurospora crassa*. Proc Natl Acad Sci 114:E7756–E7765.

36. Liu J, Wang D, Su Y, Lang K, Duan R, Wu YF, Ma F, Huang S. 2019. FairBase: a comprehensive database of fungal A-to-I RNA editing. Database (Oxford) 2019.

37. Zhu Y, Luo H, Zhang X, Song J, Sun C, Ji A, Xu J, Chen S. 2014. Abundant and selective RNA-editing events in the medicinal mushroom *Ganoderma lucidum*. Genetics 196:1047–1057.

38. Wu B, Gaskell J, Zhang J, Toapanta C, Ahrendt S, Grigoriev I V., Blanchette RA, Schilling JS, Master E, Cullen D, Hibbett DS. 2019. Evolution of substrate-specific gene expression and RNA editing in brown rot wood-decaying fungi. ISME J 13:1391–1403.

39. Kim J-S, Kwon Y-S, Bae D-W, Kwak Y-S, Kwack Y-B. 2017. Proteomic analysis of *Coprinopsis cinerea* under conditions of horizontal and perpendicular gravity. Mycobiology 45:226.

40. Kertesz-Chaloupková K, Walser PJ, Granado JD, Aebi M, Kü Es U, Kertesz-Chaloupkova K, Walser PJ, Granado JD. 1998. Blue light overrides repression of asexual sporulation by mating type genes in the basidiomycete *Coprinus cinereus*. Fungal Genet Biol 23:95–109.

41. Kuè Es U, Granado JD, Hermann R, Boulianne RP, Kertesz-Chaloupkovaâ K, Aebi M. 1998. The A mating type and blue light regulate all known differentiation processes in the basidiomycete Coprinus cinereus. Mol Gen Genet 260:81–91.

42. Moore D. 1981. Developmental genetics of *Coprinus cinereus*: Genetic evidence that carpophores and sclerotia share a common pathway of initiation. Curr Gentics 3:145– 150.

43. Swamy S, U I. 1984. Morphogenetic effects of mutations at the A and B incompatibility factors in *Coprinus cinereus*. J Gen Microbiol 130:3219–3224.

44. Thevelein JM. 1984. Regulation of trehalose mobilization in fungi. Microbiol Rev 48:42–59.

45. Jorge JA, Polizeli M de LT., Thevelein JM, Terenzi HF. 2006. Trehalases and trehalose hydrolysis in fungi. FEMS Microbiol Lett 154:165–171.

46. Domazet-Lošo T, Tautz D. 2010. A phylogenetically based transcriptome age index mirrors ontogenetic divergence patterns. Nature 468:815–818.

47. Quint M, Drost H-G, Gabel A, Ullrich KK, Bönn M, Grosse I. 2012. A transcriptomic hourglass in plant embryogenesis. Nature 490:98–101.

48. Cheng X, Hui JHL, Lee YY, Law PTW, Kwan HS. 2015. A “developmental hourglass” in fungi. Mol Biol Evol 32:1556–1566.

49. Bartel DP. 2009. MicroRNAs: target recognition and regulatory functions. Cell 136:215–233.

50. Eisenberg E, Levanon EY. 2018. A-to-I RNA editing — immune protector and transcriptome diversifier. Nat Rev Genet 19:473–490.

51. Sakamoto Y. 2018. Influences of environmental factors on fruiting body induction, development and maturation in mushroom-forming fungi. Fungal Biol Rev 32:236–248.

52. Kües U, Navarro-González M. 2015. How do Agaricomycetes shape their fruiting bodies? 1. Morphological aspects of development. Fungal Biol Rev 29:63–97.

53. Cooke WB. 1951. Ecological life history outlines for fungi. Ecology 32:736–748.

54. Fuller KK, Loros JJ, Dunlap JC. 2015. Fungal photobiology: visible light as a signal for stress, space and time. Curr Genet 61:275–288.

55. Polak E, Hermann R, Kües U, Aebi M. 1997. Asexual sporulation in *Coprinus cinereus*: Structure and development of oidiophores and oidia in an Amut Bmut homokaryon. Fungal Genet Biol 22:112–126.

56. Norros V, Karhu E, Nordén J, Vähätalo A V, Ovaskainen O. 2015. Spore sensitivity to sunlight and freezing can restrict dispersal in wood-decay fungi. Ecol Evol 5:3312–3326.

57. Rodriguez-Romero J, Hedtke M, Kastner C, Müller S, Fischer R. 2010. Fungi, hidden in soil or up in the air: light makes a difference. Annu Rev Microbiol 64:585–610.

58. Coley-Smith JR, Cooke RC. 1971. Survival and germination of fungal sclerotia. Annu Rev Phytopathol 9:65–92.

59. Waters H, Butler RD, Moore D. 1975. Structure of aerial and submerged sclerotia of *Coprinus lagopus*. New Phytol 74:199–205.

60. Golan JJ, Pringle A. 2017. Long-distance dispersal of fungi, p. 309–333. In The Fungal Kingdom. American Society of Microbiology.

61. Schreiner KM, Blair NE, Levinson W, Egerton-Warburton LM. 2014. Contribution of fungal macromolecules to soil carbon sequestration, p. 155–161. *In* Soil Carbon. Springer International Publishing, Cham.

62. Caldwell BA. 2005. Enzyme activities as a component of soil biodiversity: A review, p. 637–644. *In* Pedobiologia. Urban & Fischer.

63. Sollins P, Homann P, Caldwell BA. 1996. Stabilization and destabilization of soil organic matter: Mechanisms and controls. Geoderma 74:65–105.

64. Wallen RM, Perlin MH. 2018. An overview of the function and maintenance of sexual reproduction in dikaryotic fungi. Front Microbiol. Frontiers Media SA.

65. Heitman J. 2015. Evolution of sexual reproduction: A view from the fungal kingdom supports an evolutionary epoch with sex before sexes. Fungal Biol Rev. Elsevier.

66. Drost H-G, Gabel A, Grosse I, Quint M. 2015. Evidence for active maintenance of phylotranscriptomic hourglass patterns in animal and plant embryogenesis. Mol Biol Evol 32:1221–31.

67. Ah-Fong AMV, Kagda MS, Abrahamian M, Judelson HS. 2019. Niche-specific metabolic adaptation in biotrophic and necrotrophic oomycetes is manifested in differential use of nutrients, variation in gene content, and enzyme evolution. PLOS Pathog 15:e1007729.

68. Naranjo-Ortiz MA, Gabaldón T. 2019. Fungal evolution: major ecological adaptations and evolutionary transitions. Biol Rev 94:brv.12510.

69. Gladieux P, Ropars J, Badouin H, Branca A, Aguileta G, de Vienne DM, Rodríguez de la Vega RC, Branco S, Giraud T. 2014. Fungal evolutionary genomics provides insight into the mechanisms of adaptive divergence in eukaryotes. Mol Ecol 23:753–773.

70. Calvo AM, Cary JW. 2015. Association of fungal secondary metabolism and sclerotial biology. Front Microbiol 6:62.

71. Hoshino T, Xiao N, Tkachenko OB. 2009. Cold adaptation in the phytopathogenic fungi causing snow molds. Mycoscience 50:26–38.

72. Willetts HJ, Bullock S. 1992. Developmental biology of sclerotia. Mycol Res 96:801– 816.

73. Yablonovitch AL, Deng P, Jacobson D, Li JB. 2017. The evolution and adaptation of A-to-I RNA editing. PLOS Genet 13:e1007064.

74. Song C, Sakurai M, Shiromoto Y, Nishikura K. 2016. Functions of the RNA editing enzyme ADAR1 and their relevance to human diseases. Genes (Basel) 7:129.

75. Zhou F, Lou Q, Wang B, Xu L, Cheng C, Lu M, Sun J. 2016. Altered carbohydrates allocation by associated bacteria-fungi interactions in a bark beetle-microbe symbiosis. Sci Rep 6:20135.

76. Licht K, Jantsch MF. 2016. Rapid and dynamic transcriptome regulation by RNA editing and RNA modifications. J Cell Biol 213:15–22.

77. Wu B, Gaskell J, Held BW, Toapanta C, Vuong T, Ahrendt S, Lipzen A, Zhang J, Schilling JS, Master E, Grigoriev I V, Blanchette RA, Cullen D, Hibbett DS. 2018. Substrate-specific differential gene expression and RNA editing in the brown rot fungus *Fomitopsis pinicola*. Appl Environ Microbiol 84:e00991–18.

78. Liu H, Wang Q, He Y, Chen L, Hao C, Jiang C, Li Y, Dai Y, Kang Z, Xu JR. 2016. Genome-wide A-to-I RNA editing in fungi independent of ADAR enzymes. Genome Res 26:499–509.

79. Picardi E, Pesole G. 2013. REDItools: High-throughput RNA editing detection made easy. Bioinformatics 29:1813–1814.

80. Langmead B, Salzberg SL. 2012. Fast gapped-read alignment with Bowtie 2. Nat Methods 9:357–359.

